# Development of Loop Mediated Isothermal Amplification for Rapid and Sensitive Detection of Enteroaggregative *Escherichia coli* (EAEC)

**DOI:** 10.1101/2025.11.04.686462

**Authors:** Alazar Amare Amdiyee, Tesfaye Sisay Tessema

## Abstract

Enteroaggregative *Escherichia coli* (EAEC) is a significant etiologic agent for acute and persistent diarrhea in children and adults globally. Bacterial culture and biochemical assays are insufficient for accurately identifying EAEC strains. Currently, the HEp-2 cell assay is the gold standard method for the detection of EAEC. However, its use is restricted to the reference laboratories due to the technical complexity and infrastructure requirement. In contrast, PCR has been used as a major molecular method for detecting EAEC strains; However, its routine use in diagnostic laboratories is limited by several factors. In this study loop mediated isothermal amplification (LAMP) was developed for affordable, rapid and specific detection of EAEC. The assay was developed by using a specifically designed lamp primer targeting aaic genes to enable precise detection of EAEC. The performance of the developed assay was evaluated by using 60 locally isolated bacterial strains. The assay exhibited 100% sensitivity and a specificity. The developed LAMP assay could detect up to 0.098pg of DNA/reaction. In contrast, the conventional PCR exhibited a detection limit of 0.98pg/reaction. In pure culture, the developed LAMP assay has a detection limit of 80 CFU/mL, which is higher than 8× 10^2^ CFU/mL for conventional PCR, indicating the LAMP assay’s 10-fold higher sensitivity than conventional PCR. Furthermore, in spiked stool sample the lowest detection limit for LAMP was 8 × 10^2^ cfu/ g stool. In contrast, 8 × 10^4^ cfu/g stool for PCR. Notably, the LAMP assay shows 100-fold higher sensitivity than conventional PCR.

## 1. Introduction

Enteroaggregative *Escherichia coli* (EAEC) is a widespread diarrheagenic *E. coli* (DEC) pathotypes associated with episodes of acute and persistent diarrhea in children and adults worldwide. It accounts for a considerable portion of the global diarrheal disease burden, being implicated in both epidemic and endemic cases [1–3]. A meta-analysis of 41 studies shows EAEC as a significant cause of acute diarrheal illness in developing countries[4]. The pathophysiology of EAEC strains is mediated by their interaction with human and animal intestinal cells. The in vitro, vivo, and ex vivo investigations show EAEC can stick to the jejunal, ileal, and colonic epithelium and exhibits a distinctive ‘stacked-brick’ adherence pattern, which creates a robust biofilm within a mucus layer, followed by cytotoxic and proinflammatory effects[5]. The clinical symptoms of EAEC infection comprises watery diarrhea, occasionally with blood and mucus [6].

Bacterial culture and biochemical tests are inadequate for reliable identification of EAEC strain as they exhibit similar phenotypic characteristics to normal flora and other pathogenic *E. coli* strains. The gold standard method for identifying EAEC is a HEp-2 cell culture-based method, which detects aggregative adherence (AA) pattern exhibited by the bacteria on epithelial cells[7–9]. However, the implementation of HEp-2 cell assay is limited due to the requirement of cell culture facilities, which available only in reference laboratories and prolonged processing time[10, 11].In contrast, Polymerase chain reaction (PCR) has been used as a major molecular approach for detecting EAEC strains based on the presence/absence of specific virulence genes, including aaic (aggR-activated Island C) located on conserved chromosomal region of EAEC, and encodes type VI secretion system[12, 13]. Despite PCR’s more applicability than HEp-2 cells assay, it has certain limitation including intensive sample preparation in order to eliminate amplification inhibitors, difficulty in optimizing reaction condition, need for skilled personnel and reliance on expensive thermal cycler equipment, which limits the applicability in routine laboratories. In contrast, loop-mediated isothermal amplification (LAMP) was introduced as alternative to PCR, offering a simple, rapid, and cost-effective detection of the target pathogen [14, 15].

Loop-mediated isothermal amplification (LAMP) is first introduced by Notomi et al. in 2000. In contrast to PCR, LAMP is performed under a constant temperature by using *Bacillus stearothermophilus* (Bst) DNA polymerase, which facilitates auto-cycling strand displacement DNA synthesis. LAMP employs a set of four to six specifically designed primers, thereby significantly enhance both the sensitivity and specificity of the assay [16–18]. LAMP requires affordable equipment, such as a water bath or heat block that maintains a constant temperature, making it readily adaptable for use in routine laboratories[19, 20]. Recently, loop-mediated isothermal amplification (LAMP) has been effectively used in detection of various pathogens, including *Salmonella Typhi, COVID-19*, *Ebola virus*, *Staphylococcus aureus* and Malaria parasites [21–25]. Despite the widespread prevalence of Enteroaggregative *Escherichia coli* (EAEC) as a bacterial cause of diarrhea, it remains a critical need for the development of rapid diagnostics such as Loop-Mediated Isothermal Amplification (LAMP) assay, that enables rapid, affordable, and user-friendly target detection. Therefore, the objective of this study was to develop a LAMP-based diagnostic method for effective identification of Enteroaggregative *Escherichia coli* (EAEC) strains.

## 2. Materials and Methods

### 2.1. Bacterial strains

In this study a total of 60 bacterial strains were used for the development and evaluation of the LAMP assay. These include 10 EAEC, 10 ETEC,10 EPEC, ,10 STEC,4 EIEC, 2 EHEC,10 non photogenic *E. coli* strains & 4 non-*E. coli* bacterial species (*Pseudomonas, Klebsiella pneumoniae*, *Salmonella spp*, and *Staphylococcus aureus*). The number of EHEC and EIEC is low due to few isolates available in our culture collection and inaccessibility of getting more isolates. All strain were preserved in glycerol stocks at Prof. Tesfaye Sisay Tessema’s laboratory in the Institute of biotechnology, Addis Ababa university. *E. coli* strains were cultured in eosin methylene blue (EMB) agar, while non-*E. coli* bacterial strains were cultured on nutrient agar. All bacterial cultures were incubated aerobically at 37°C for 24hrs.The bacterial isolates were sub cultured in 10ml of tryptone soya broth (TSB) and incubated for 24 hrs at 37 °C.

### 2.2. DNA Extraction

One ml of each bacterial culture grown in TSB was transferred into 1.5ml sterile Eppendorf tube and centrifuged at 15000rpm for 10 min. The supernatant was discarded and the resulting pellet was resuspended in 200 µL of nuclease free water. A centrifugation at 15000rpm for 10 min was followed. Upon discarding the supernatant, 100 μL of TE buffer was added to the pellet and incubated at 100C° for 10min.The sample was then centrifuged at 10, 000 rpm for 2 min and cooled on ice for 1min. The resulting supernatant was transferred to new sterile Eppendorf tube and stored at −20C° for uses as the DNA template in LAMP and PCR assays.

### 2.3. PCR Amplification

The conventional PCR was conducted in parallel to confirm the LAMP assay’s result accuracy by utilizing specific PCR primers provided in Table 1. Each PCR assay was carried out in 25 µl reaction volume containing, 2.5 µL of MgSO_4,_ 2.5 µL PCR buffer, 0.5 µL of each forward and reverse primers, 0.5 µL of dNTP, 8U of Taq DNA polymerase,15 µL of nuclease free water and 3 µL of DNA template. The same reaction mixture without template DNA was used as a negative control. Amplification was performed with an initial denaturation temperature of 95 °C for 4 min followed by 37 amplification cycles each of them consisting a 30 s denaturation, 30 s annealing and 1 min extension. The last amplification cycle included final extension at 72 °C for 10min. All amplifications were carried out in A300 Thermal Cycler (Long gene Scientific Instruments, China). The PCR product was subjected to 1.5% agarose gel electrophoresis to analyze DNA fragments. Electrophoresis was carried out at 100 V for 1hr in 1X TAE buffer. A 100-bp DNA ladder was used to estimate the product size and a UV gel documentation system were used to visualize and record the resulting DNA bands.

**Table 1:**
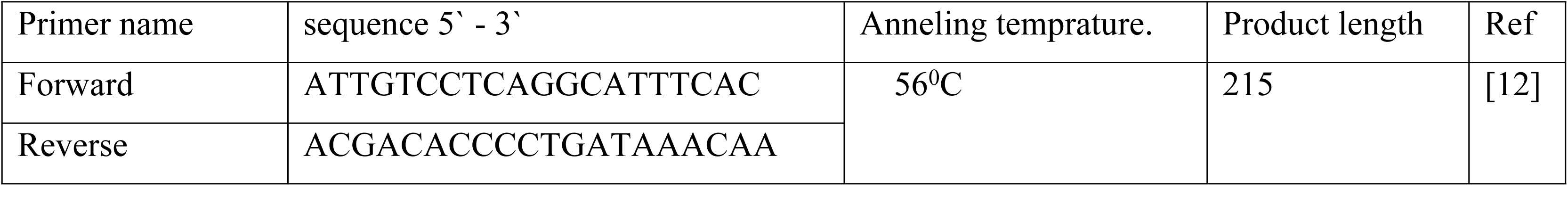
Nucleotide sequences and amplification features of PCR primers.

### 2.4. LAMP primer designing

LAMP primers were designed by using the following software: https://www.optigene.co.uk/custom services lamp designer software, https://lamp.neb.com/#!/,targeting aaic (Gene bank accession no: >NC_017626.1(4895043-4895549)).

### 2.5. LAMP optimization and detection of amplification products

A single-factor optimization approach was applied to establish optimal LAMP reaction condition based on the initial 25 µL reaction mixture containing: 2.5 µL of Isothermal amplification buffer ,1 µL of Bst DNA polymerase (8 U, New England Biolabs), 10.0 mM MgSO₄, 1.2 mM dNTPs, 1.6 µM each of FIP and BIP primers, 0.2 µM each of F3 and B3 primers, 0.4 µM each of LF and LB primers, and 2.0 µL of template DNA. Sterile double-deionized water (ddH₂O) was used as a negative control in place of template DNA. The mixture was incubated at 65°C for 60 minutes and terminated by heating at 80 °C for 5 minutes. Optimization was carried out by independently varying five critical reaction components including; temperature, MgSO₄ concentration, dNTP concentration, Bst DNA polymerase concentration and incubation time, each of the five Factor was tested at multiple levels to assess their impact on amplification efficiency.

The amplification product was detected by using a combination of methods including: visual turbidity evaluation, SYBR Green I staining, and agarose gel electrophoresis. SYBR Green I detection was carried out by adding 1µL of SYBR Green I to the reaction mixture, and the color in the mixture was observed by naked eye. In the Prescence of amplified DNA products the SYBR green turned to green color, while the reaction without DNA amplification remained orange. For the gel electrophoresis analysis, 3 µL of each reaction product was subjected to 1.5% agarose gel stained with ethidium bromide; the electrophoresis was carried out by 100 V for 1hr in 1X TAE buffer. In the presence of target amplification, a ladder of DNA bands was visualized and documented by using UV transilluminator gel documentation system.

### 2.6. Sensitivity, Specificity and Detection limit of the LAMP assay

The diagnostic performance of the developed LAMP assay was evaluated by using 60 bacterial isolates, including 10 Enteroaggregative *Escherichia coli* (EAEC) strains and 50 non-EAEC bacterial isolates. Sensitivity was defined as the proportion of EAEC strains correctly identified as positive, while specificity referred to the proportion of non-EAEC strains accurately classified as negative. These criteria were used to assess the assay’s capability in detecting EAEC under defined experimental conditions. For determining lower detection limit of the developed LAMP assay, overnight broth cultures of *E. coli* strain was serially diluted from 10^−1^ up to 10 ^−10^. Briefly, a single colony of each strain was inoculated separately into 10 ml of fresh TSB and incubated at 37°C for 24hr. The cultures were diluted 10-fold serially in normal saline, then an aliquot of 1ml of each dilution was used for DNA template preparation and another 1 ml plated on plate count agar for CFU count to evaluate the detection limit of the assay in terms of lowest numbers of cells that could be detected. For determining lower number of DNA concentration/reaction the extracted DNA from original culture was subjected to 10-fold serial dilutions ranging from 10⁻¹ to 10⁻¹⁰. Two µL of template DNA from each dilution was used for LAMP and PCR assay. Finally, the lower detection limit of the assay was presented as in terms of lower number of DNA concentration/reaction and numbers of CFU/ml.

### 2.7. LAMP Application on Human Fecal Samples Spiked With EAEC

One g fecal sample was diluted in 9 ml of phosphate-buffered saline (PBS). Samples were vortexed for 10s, allowed to settle for 2 min, vortexed again; then, 1ml of stool suspension was added to 10 separate tubes and spiked with 1 ml of serially diluted (10^9^ to 10°) EAEC bacterial suspension. Afterwards 1ml of each spiked fecal sample dilution was transferred to an Eppendorf tube containing 10X EDTA. Samples were vortexed for 30 s and then centrifuged at 15000 rpm for 15 min; the supernatants were discarded, and 300 μl of nuclease free water was added; then centrifuged again for 15000 rpm for 10 min. The supernatants were discarded; then the pellets were suspended in 100 μl 1x TE buffer and boiled at 100°C for 15 min. After boiling centrifugation was performed at 15,000 rpm for 5min. Then the supernatant, to be used as template DNA was transferred to a new sterile Eppendorf tube; then the template subjected to PCR and LAMP assay. The feces without EAEC was used as a negative control.

### 2.8. Data Analysis

The key diagnostic performance parameters, including specificity, sensitivity, positive predictive value (PPV), negative predictive value (NPV), and accuracy were calculated using a 2×2 contingency table and analyzed by MedCalc. software version 23.11.3. Additionally, Cohen’s kappa (κ) statistic was employed to measure the level of agreement between the PCR and LAMP assay results.

## 3. Results

### 3.1. Primer design for LAMP assay

In this study a set of 4 lamp primers were designed by targeting aaic (Gene bank accession no: >NC_017626.1(4895043-4895549)) using the following software: https// www.optigene.co.uk/custom services lamp designer software, http: lamp.neb.com. The designed primer includes inner primers (Forward inner primer-FIP and Backward inner primer-BIP) and outer primers (Forward outer primer-F3 and Backward outer primer-B3). The overall information of primer sequences is provided in (Table 2).

**Table 2:**
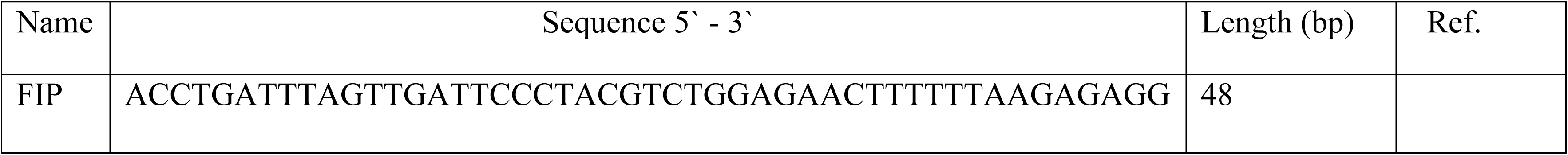
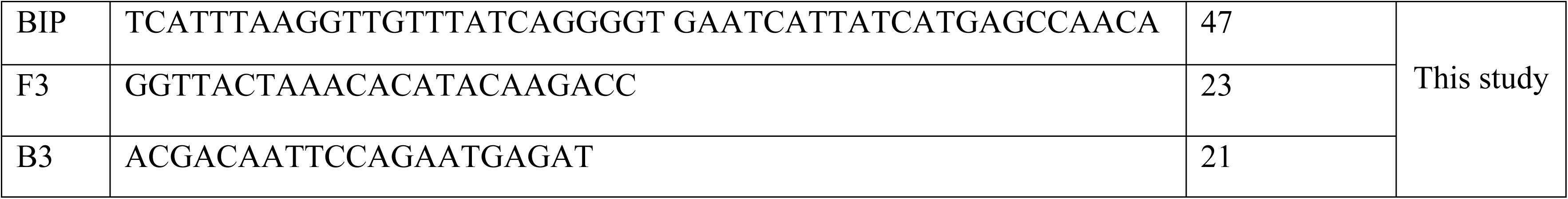
Information of designed LAMP primer sequences.

### 3.2. Optimization process of LAMP assay

Initial re-confirmation of EAEC strain was carried out by PCR (Figure 1a). Subsequently, the LAMP assay was performed to verify the designed lamp primer effectiveness based on the initial reaction condition as described in the methods section. The amplification result was evaluated by using: turbidity assessment (Figure 1b) agarose gel electrophoresis (Figure 1c) and SYBR Green I dye detection (Figure 1d). Optimal reaction condition of the LAMP assay was established after systematic optimization of five critical factors. This stepwise optimization process allowed the identification of the most effective combination of parameters to enhance reaction efficiency and specificity of the LAMP assay. The detailed optimization process and corresponding results for each factor are presented below.

**Figure 1:**
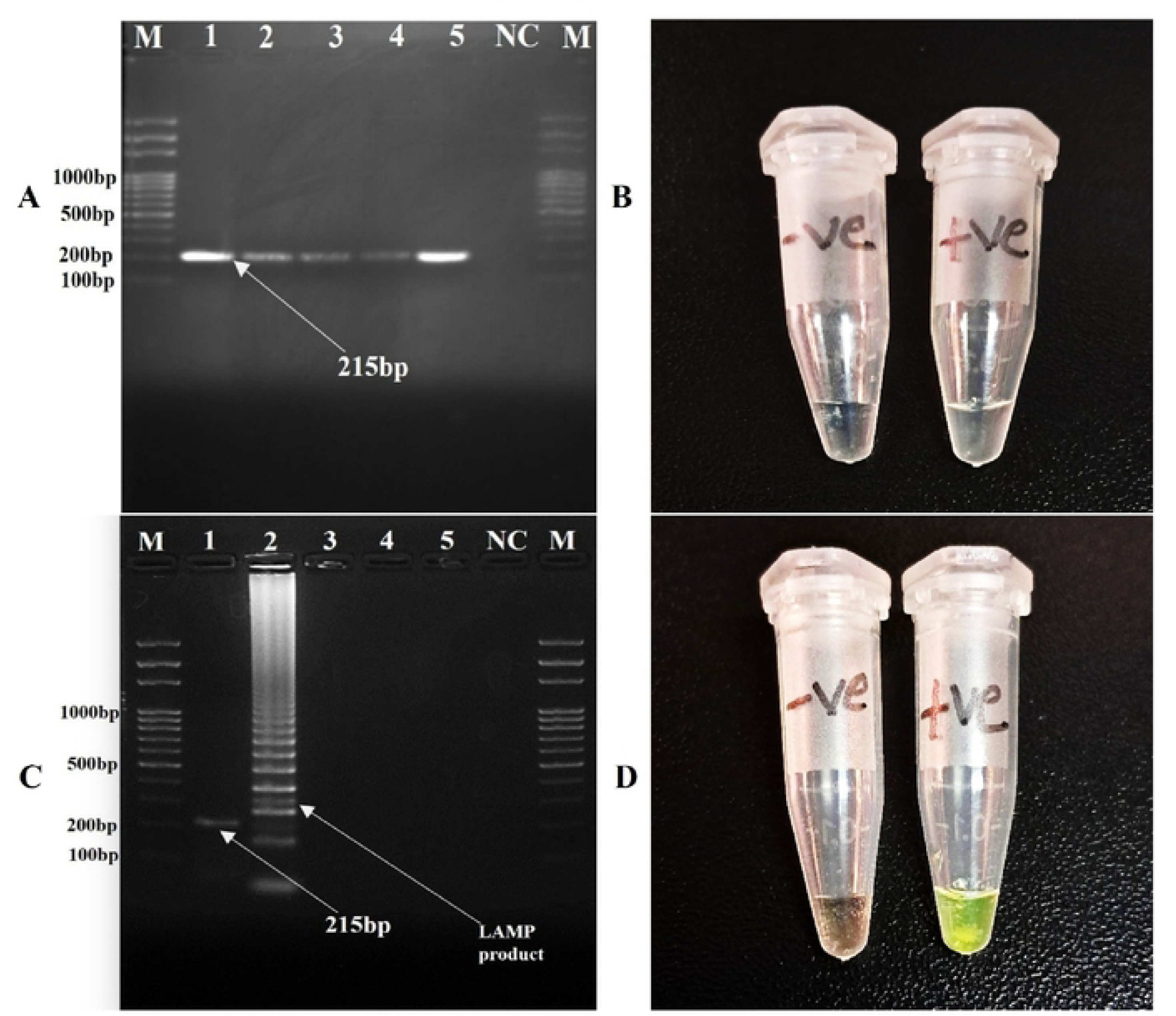
PCR and LAMP based Detection of EAEC. A) Shows agarose gel electrophoresis PCR product: Lane M=100 bp marker, lane 1= 024, Lane2: 61C, Lane 3: 72I, lane 4:7I, lane 5: 73I, lane 6: NC.7B) visual turbidity assessment of LAMP Product. C) Agarose gel electrophoresis: Lane M=100 bp marker, lane 1= PCR product of EAEC 215bp, Lane 2= EAEC LAMP product, Lane 3-NC=negative control for LAMP and PCR D) SYBR Green I detection of LAMP product.

#### 3.1.1. Optimization of MgSO4 Concentration

The optimization of MgSO4 concentration was performed by testing different concentration of MgSO4 ranging from 2.0 to 18 mM and the result showed a green color in a range from 4.0 to 18.0 mM after adding of SYBR green I. The green color in SYBR Green I staining indicates the presences of amplification at tested concentration and the appearance of orange color at 2.0 mM concentration indicates the absence of amplification (Figure 2a). In agarose gel electrophoresis, the LAMP products appeared as a ladder-like bands from the reaction with the MgSO4 concentrations ranging from 4.0 to 18.0 mM. A robust band was appeared at 10.0 mM. Therefore,10.0 mM MgSO4 concentration for the reaction was considered as an optimal concentration (Figure 2b).

**Figure 2:**
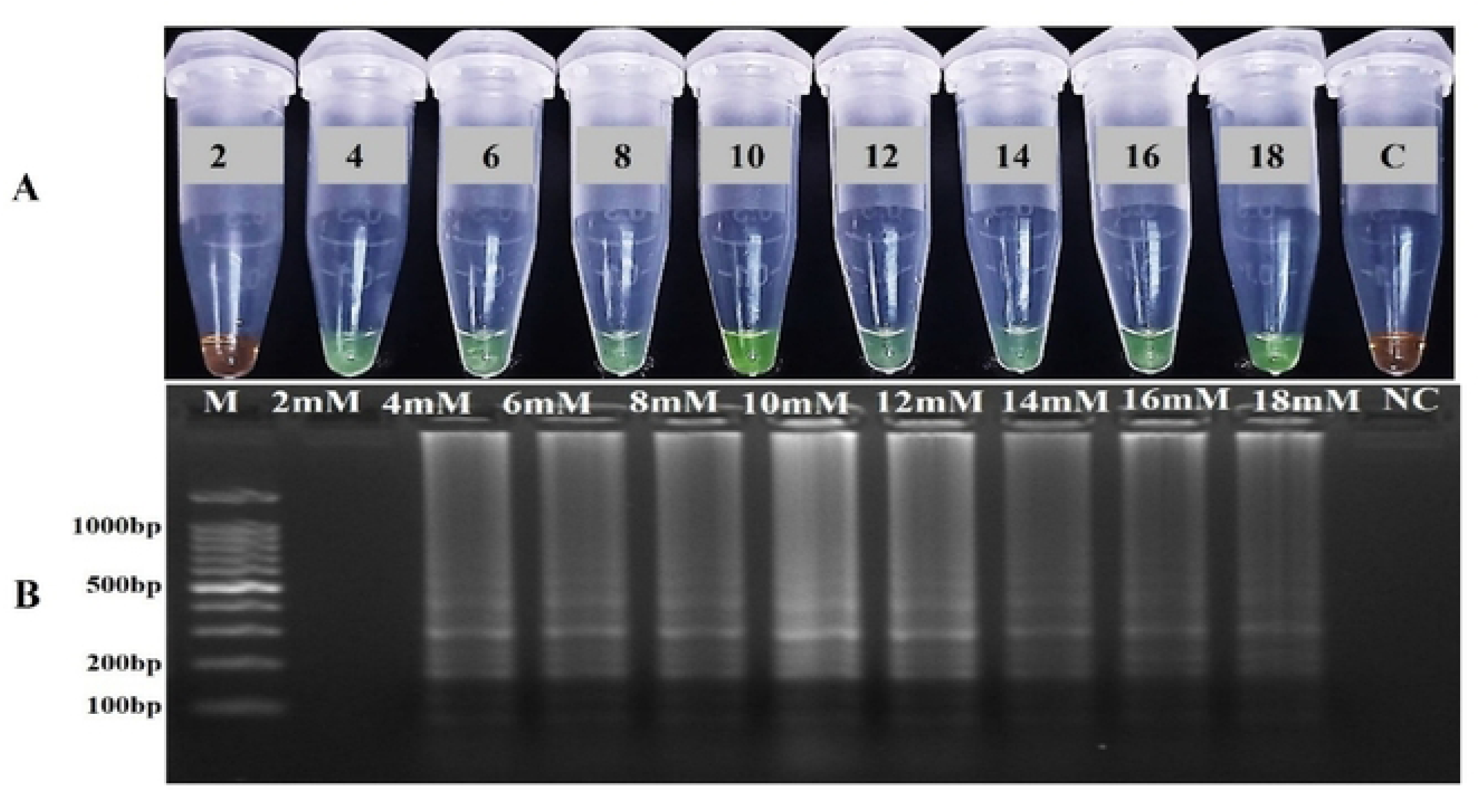
LAMP reactions with different concentrations ranging from 2.0 to 18 mm of MgSO4 solution (A). Assessments of LAMP products based on SYBR Green I visualization (B). Agarose gel electrophoresis result.

#### 3.1.2. Optimization of Reaction Temperature

Different temperatures ranging from 57°C to 65°C were evaluated; a fluorescent green color was displayed at all temperature ranges after SYBR Green I staining (Figure 3a) and in the agarose gel a ladder-like bands were observed in all evaluated reaction temperatures. The brightest band was observed at 61°C (Figure 3b). Therefore, 61°C considered as an optimal temperature for the LAMP reaction.

**Figure 3:**
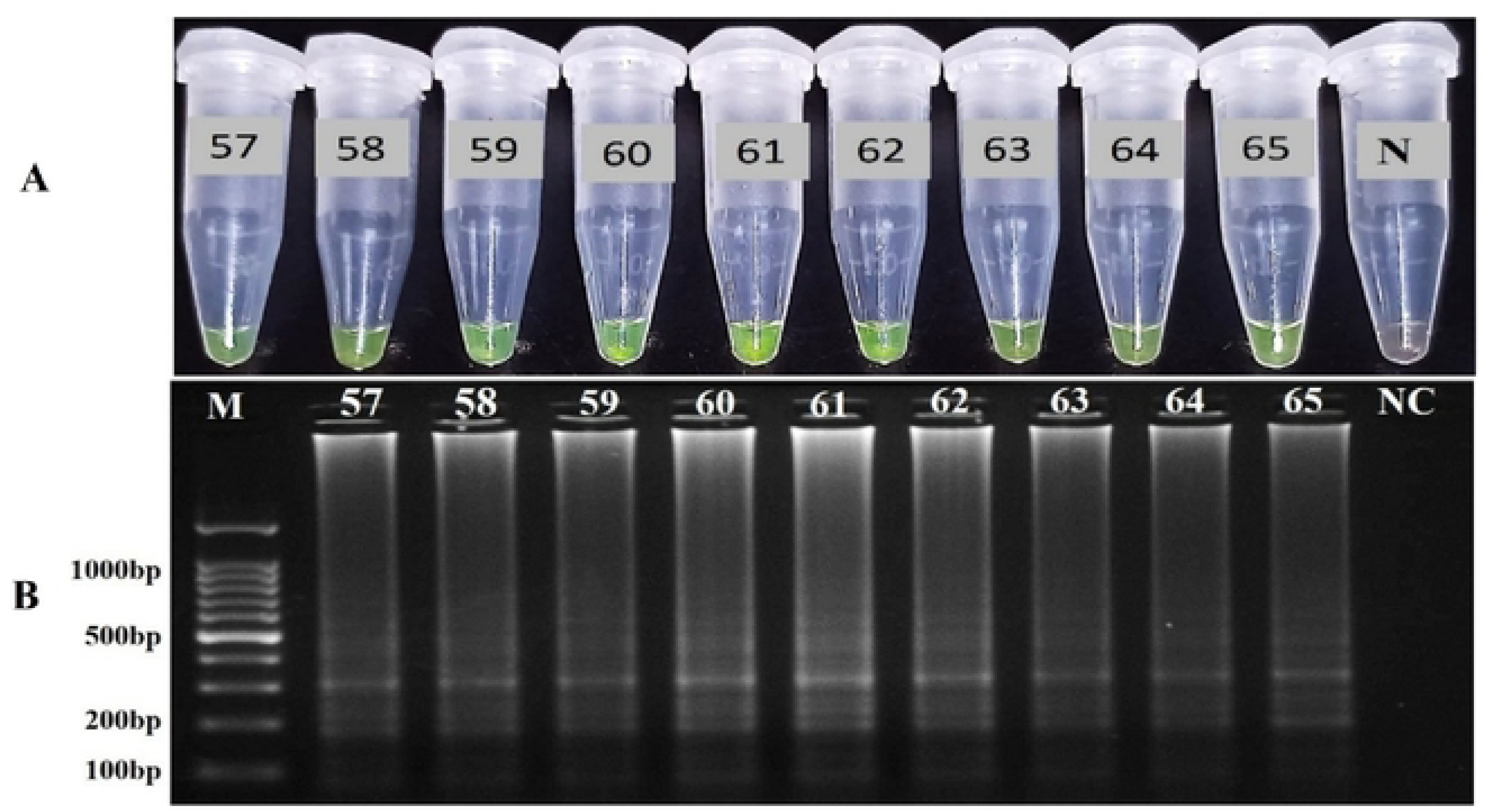
Results of the LAMP reactions at a temperature of 57,58,59,60,61,62,63,64,65. A) SYBR green I detection. B) Agarose gel electrophoresis results.

#### 3.2.3. Optimization of incubation Time

To determine optimal time for LAMP reaction the incubation time ranging from 10 up to 70 min was evaluated. No color change was observed by SYBR green I staining until 40 min, as the color remained orange, starting from 45 up to 65 min a fluorescent green color was observed indicating positive result and at 70min no color change was observed (Figure 4a). In agarose gel electrophoresis, the LAMP products displayed ladder-like bands starting from 40 up to 65 min. No ladder-like bands were observed between 10 up to 40 and 70 min. A robust band was produced at 60min.Therefore, the 60min incubation time for LAMP reaction was considered as an optimal temperature for the LAMP reaction (Figure 4b).

**Figure 4:**
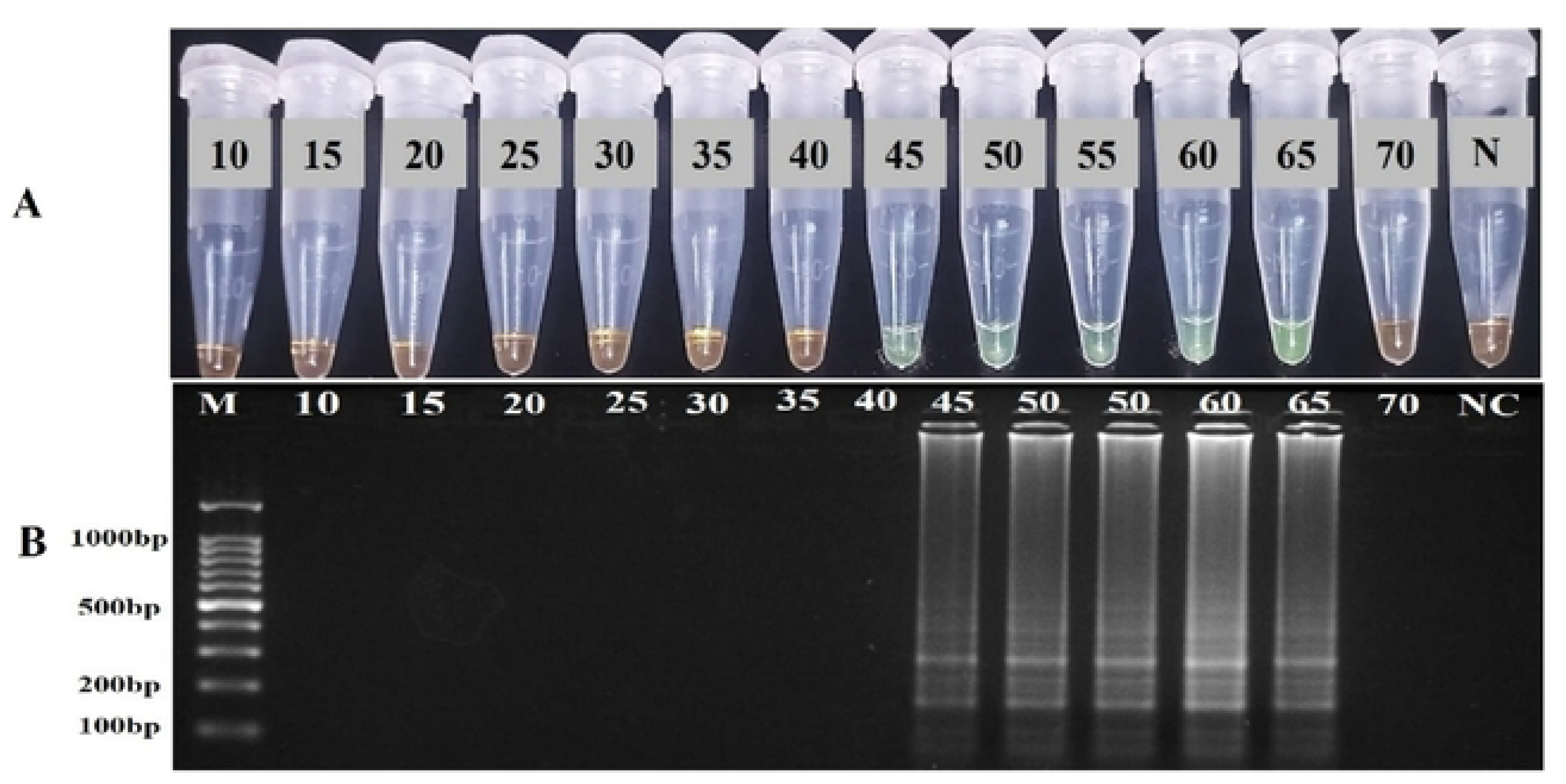
Profiles of LAMP reactions with different reaction times. Assessments of LAMP products based on SYBR Green I visualization of color change tubes 1 to 14 respectively, LAMP reaction with reaction time of 10, 15, 20, 25, 30, 35,40,45,50, 55,60,65, and 70 min, respectively; tube 14, negative control with ddH2O for 60 min.

#### 3.2.4. Optimization of Bst polymerase Concentration

The bst polymerase concentration ranging from 2-16U were evaluated and a fluorescent green color was displayed in all evaluated concentration after SYBR Green I staining, indicating the presence of amplification (Figure 5a). A robust band was displayed at 8.0 U of bst DNA polymerase. Therefore, 8U bst polymerase considered as optimal concentration for the reaction (Figure 5b).

**Figure 5:**
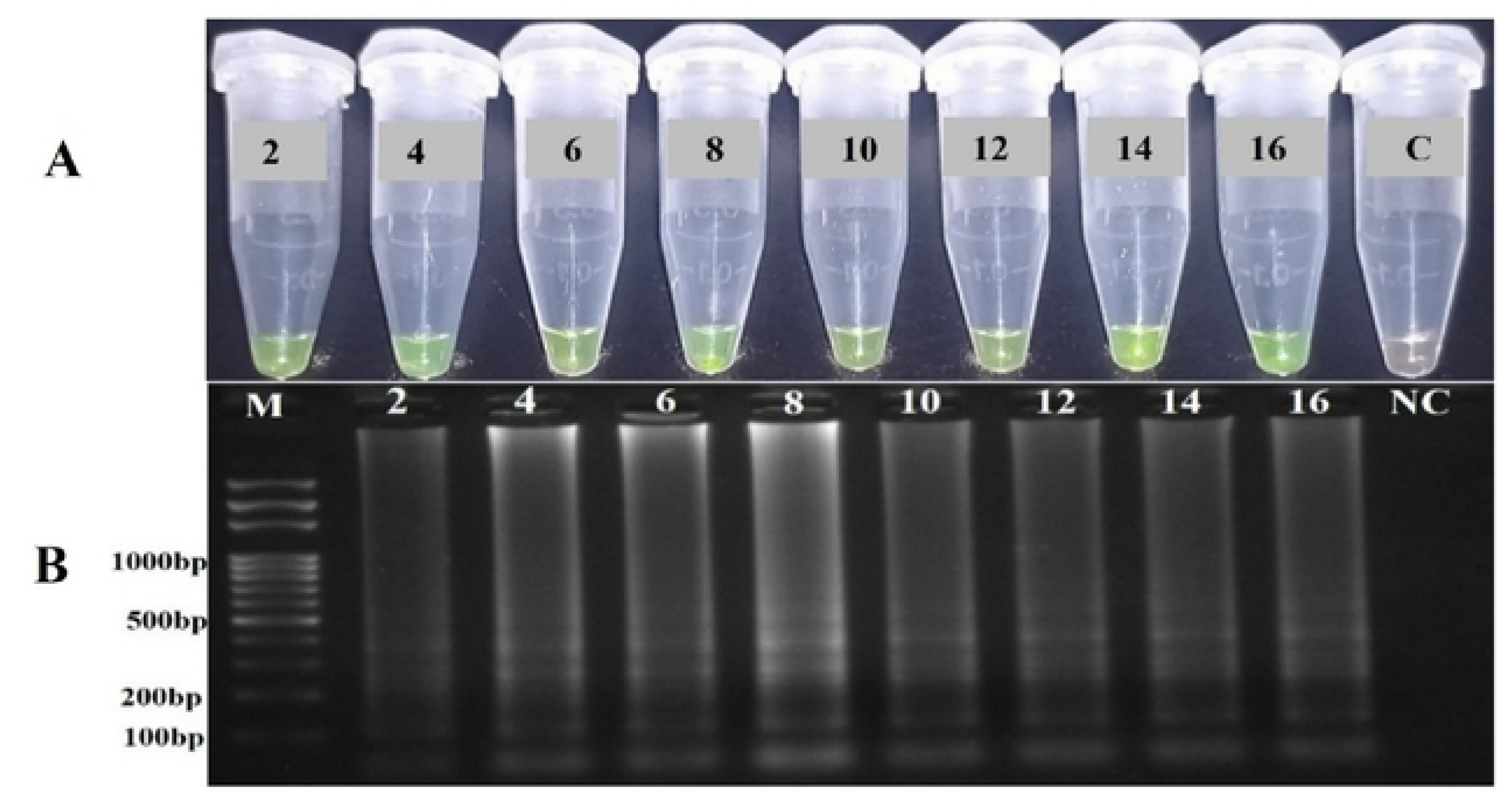
Result of LAMP reactions with different Bst polymerase concentrations, 2.0, 4.0, 6.0, 8.0, 10.0, 12.0, 14.0 ,16.0U, respectively. A) SYBR Green I visualization. (B) Agarose gel electrophoresis result.

#### 3.2.5. Optimization of DNTPs Concentration

All nine different concentration of dNTPs ranging from 0.6 −2 mM showed a fluorescent green color in SYBR Green I staining, indicating the target DNA amplified with dNTP concentrations ranging between 0.6 and 2 mM (Figure 6a). Agarose gel electrophoresis result showed a ladder-like bands in all evaluated dNTPs concentration, LAMP products showed a robust band in reactions with 1.4mM dNTPs. Therefore, 1.4mM considered as an optimal dNTPs concentration for the LAMP reaction (Figure 6b).

**Figure 6:**
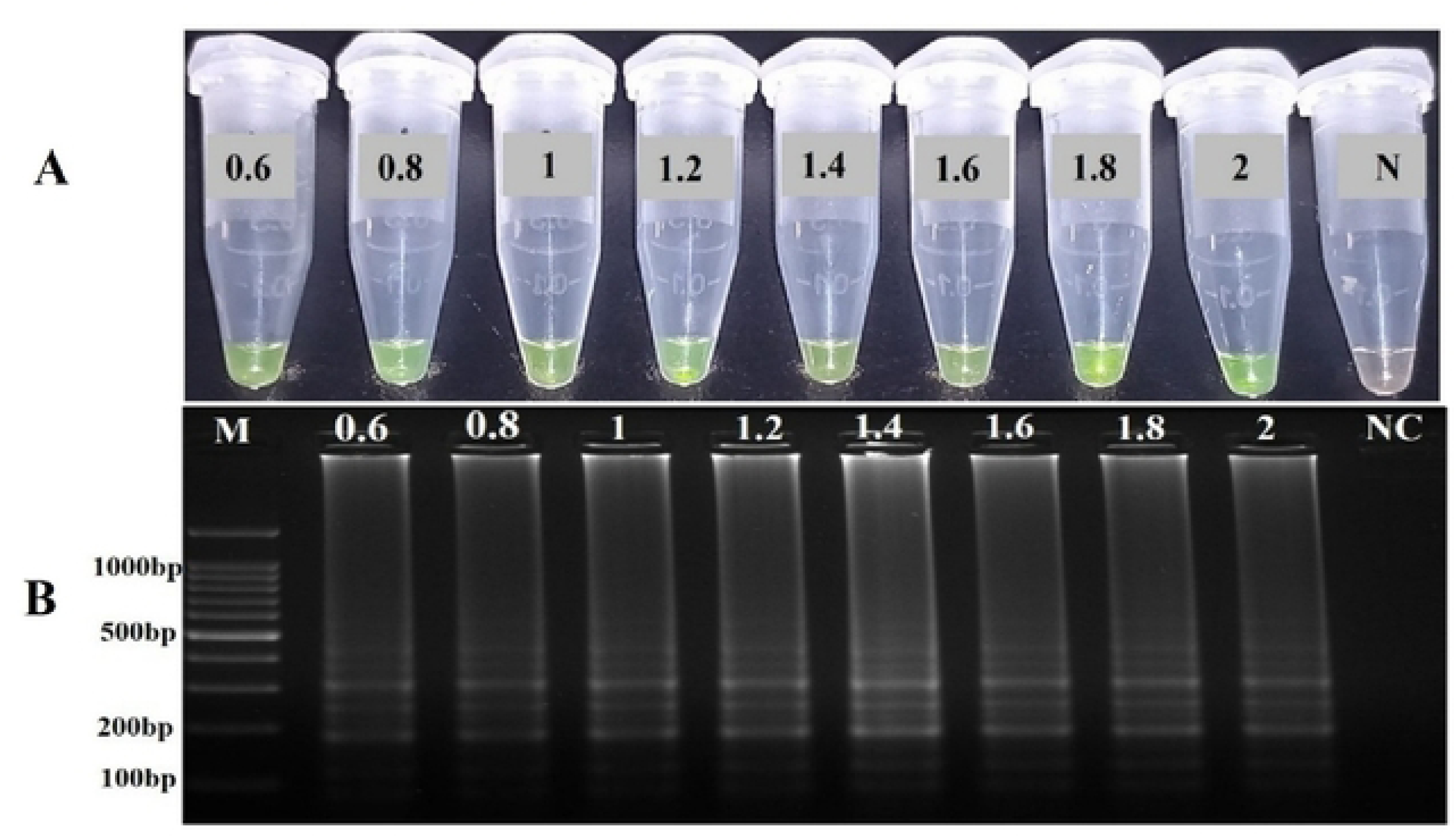
Result of LAMP reactions with different dNTPS concentrations ranging from 0.6, 0.8, 1.0, 1.2, 1.4 ,1.6,1.8,2 respectively. A) Shows SYBR Green I visualization. B) Shows agarose gel electrophoresis result.

#### 3.2.6. Application of Optimized LAMP Protocol

The optimum reaction conditions were established after evaluations of five critical reaction factors, the final lamp reaction was carried out in a total of 25 µL volume containing 2.5 µL of isothermal amplification buffer, 10 mM of MgSO4 solution, 1.4 mM of dNTPs, 1.2 mM each of FIP and BIP, 0.4 mM each of F3 and B3, 0.2 mM of LB, 8.0 U of Bst polymerase and 2.0 µL of the target DNA template and the reaction mixture was incubated at 61 C° for 60 min. This optimized condition was used for subsequent LAMP reaction on ten EAEC strain. The result of LAMP amplification was verified by SYBR Green I staining color change, and agarose gel electrophoresis, which revealed the characteristic of ladder-like bands of LAMP products (Figure 7).

**Figure 7:**
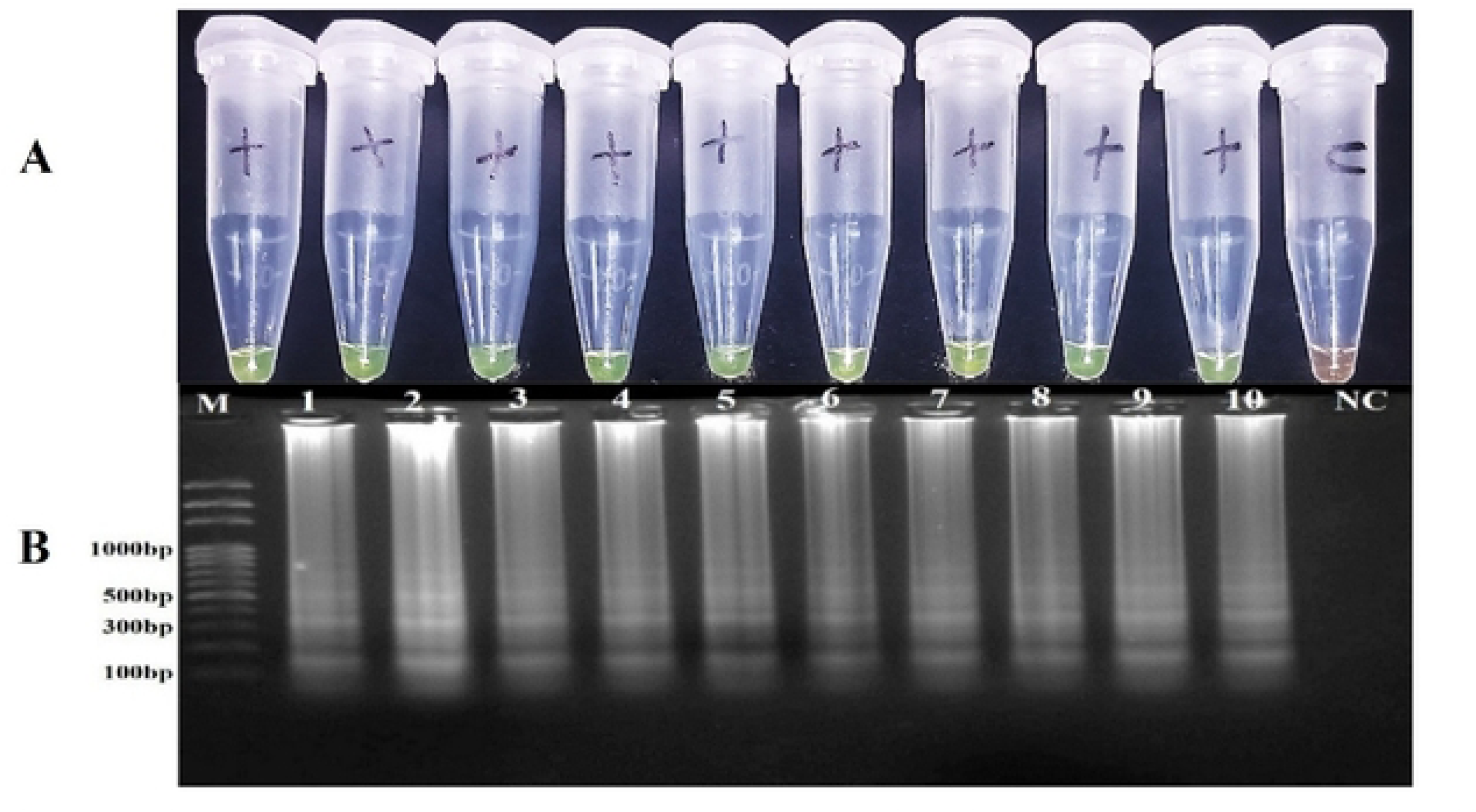
SYBR green I detection of Lamp product of EAEC amplification; Tube 1-10 shows a positive result: Tube 1: 024, Tube 2: 61C, Tube 3: 72I, Tube 4:7I, Tube 5:73I, Tube 6:72E, Tube 7: 82F, Tube 8: EV10, Tube 10: EV71, Tube11: IH 64, Tube 11: negative control (ddwater). B) Gel electrophoresis detection of LAMP product of EAEC isolates shows ladder like appearance; lane M: DNA ladder (100plus), lane 1: 024, lane 2: 61C, lane3: 72I, lane 4: 7I, lane 5: 73I, lane 6: 72E, Lane 7: 82F, lane 8: EV10., lane 9: EV71, lane 10: IH 64, lane 11: negative control (ddwater).

### 3.3. Validation of LAMP assay

PCR was performed in parallel with LAMP to validate the result of 60 bacterial isolates tested by the LAMP assay (Table 3). Based on PCR finding, the test isolates were classified as true positive when it was EAEC; whereas, the test isolates were declared as true negative when it was not EAEC. In contrast, test positive means that the isolate was positive in the LAMP test and test negative means that the isolate was negative in the LAMP test. To measure the sensitivity, specificity and the overall performance of the LAMP assay (2×2 contingency matrix) was used (Table 4).

**Table 3:**
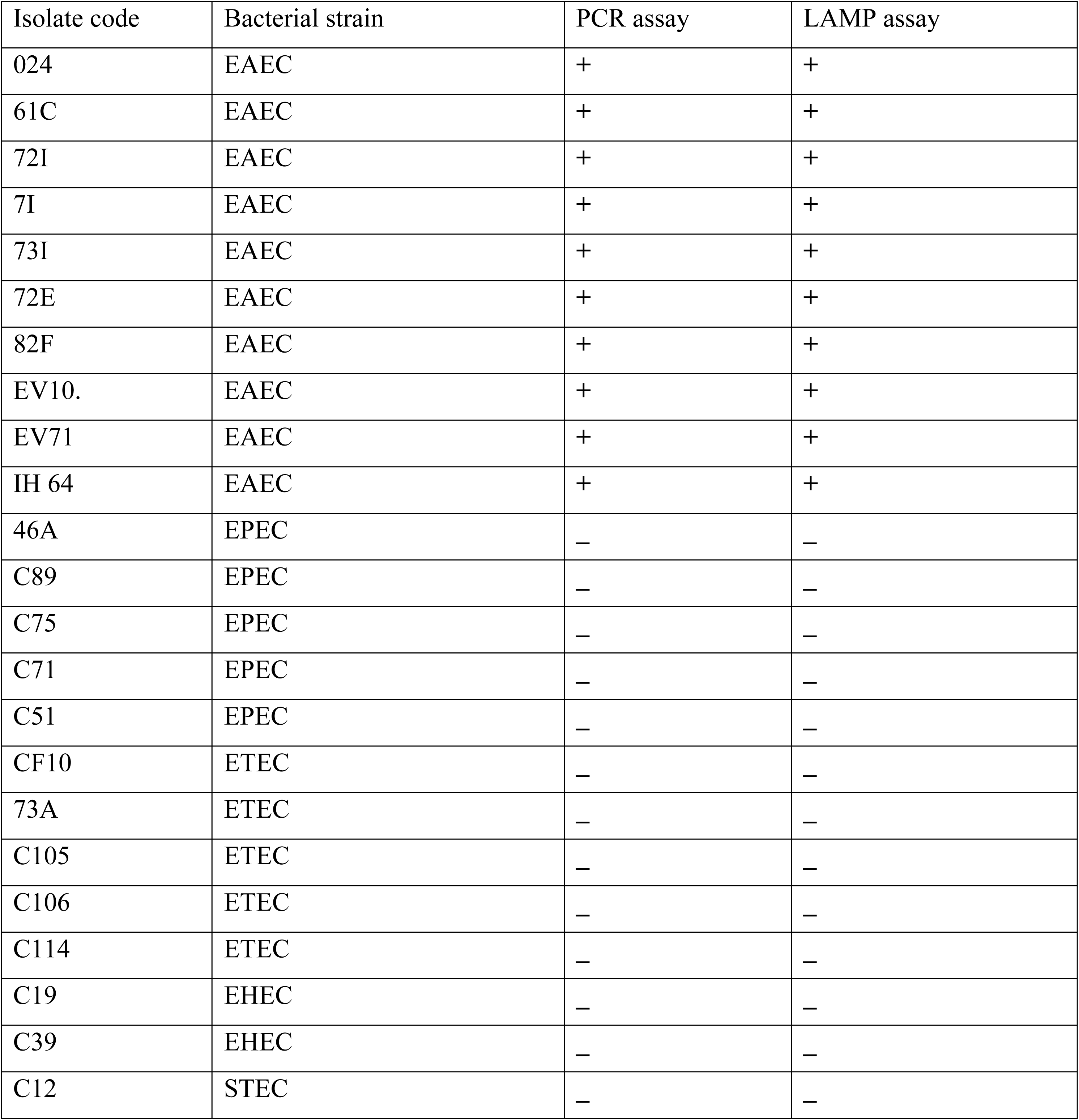
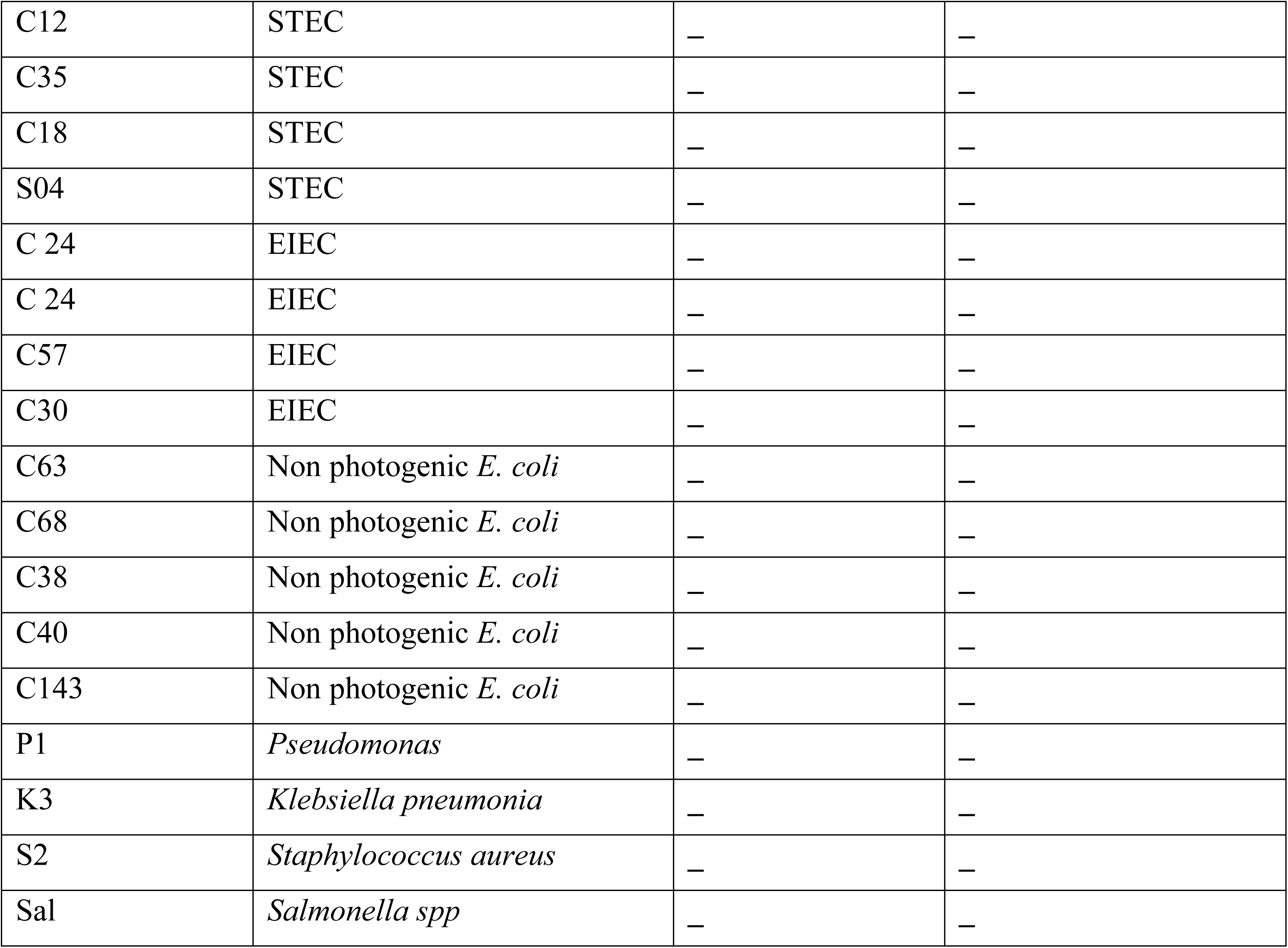
Parallel Comparison of LAMP and PCR.

| Isolate code | Bacterial strain | PCR assay | LAMP assay |
| --- | --- | --- | --- |
| 024 | EAEC | + | + |
| 61C | EAEC | + | + |
| 72I | EAEC | + | + |
| 7I | EAEC | + | + |
| 73I | EAEC | + | + |
| 72E | EAEC | + | + |
| 82F | EAEC | + | + |
| EV10. | EAEC | + | + |
| EV71 | EAEC | + | + |
| IH 64 | EAEC | + | + |
| 46A | EPEC | – | – |
| C89 | EPEC | – | – |
| C75 | EPEC | – | – |
| C71 | EPEC | – | – |
| C51 | EPEC | – | – |
| CF10 | ETEC | – | – |
| 73A | ETEC | – | – |
| C105 | ETEC | – | – |
| C106 | ETEC | – | – |
| C114 | ETEC | – | – |
| C19 | EHEC | – | – |
| C39 | EHEC | – | – |
| C12 | STEC | – | – |
| C12 | STEC | – | – |
| C35 | STEC | – | – |
| C18 | STEC | – | – |
| S04 | STEC | – | – |
| C 24 | EIEC | – | – |
| C 24 | EIEC | – | – |
| C57 | EIEC | – | – |
| C30 | EIEC | – | – |
| C63 | Non photogenic *E. coli* | – | – |
| C68 | Non photogenic *E. coli* | – | – |
| C38 | Non photogenic *E. coli* | – | – |
| C40 | Non photogenic *E. coli* | – | – |
| C143 | Non photogenic *E. coli* | – | – |
| P1 | *Pseudomonas* | – | – |
| K3 | *Klebsiella pneumonia* | – | – |
| S2 | *Staphylococcus aureus* | – | – |
| Sal | *Salmonella spp* | – | – |

**Table 4:** 2×2Contingency matrix table generated by MedCalc Version 23.2.1 Software.

| Test | Present | No | Absent | No | Total |
| --- | --- | --- | --- | --- | --- |
| Positive | TP | 10 | FP | 0 | 10 |
| Negative | FN | 0 | TN | 50 | 50 |
| Total | - | 10 |  | 50 | 60 |

#### 3.3.1. Specificity of the LAMP assay

The developed LAMP assay specificity was evaluated by using 60 bacterial isolates containing 10 EAEC, 46 non-EAEC *Escherichia coli* strains and 4 non-*E. coli* bacterial species. As shown in (Figure 8) no positive result was observed among non-EAEC *E. coli* isolates. In addition, no amplification was observed in the non-*E. coli* species, confirming the assay’s genus-level specificity and absence of cross-reactivity (Figure 9). In addition, The LAMP assay yielded negative result for all 50 non -EAEC bacterial, indicates absence of false-positive results confirms the assay’s 100% analytical specificity (Table 5).

**Figure 8:**
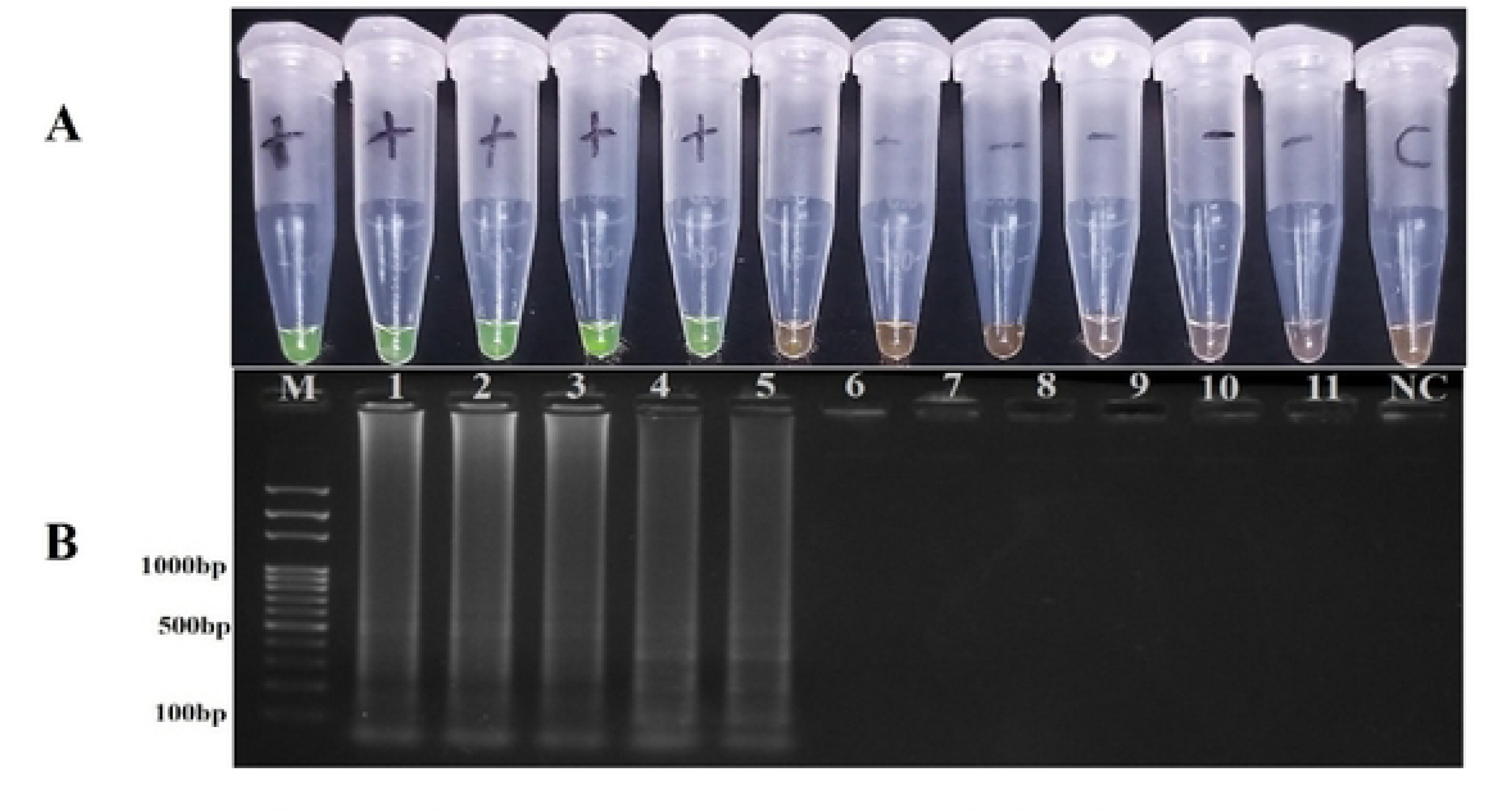
Specificity of the LAMP assay on non EAEC *E. coli***; A**) SYBR green I detection: Tube 1: 024(EAEC), Tube 2: 61C(EAEC), Tube 3: 72I(EAEC), Tube 4: 7I(EAEC), Tube 5: 73I(EAEC), Tube 6:C19(EHEC), Tube 7: 46A (EPEC), Tube 8: 73A(ETEC), Tube 9:C24(EIEC), Tube 10:C12 (STEC), Tube 11: Non photogenic E. coli, lane 12: negative control (ddwater). **B)** Agarose gel electrophoresis result: lane M: DNA ladder (100plus), lane 1: 024(EAEC), lane 2(EAEC): 61C(EAEC), lane3: 72I(EAEC), lane 4: 7I(EAEC), lane 5: 73I(EAEC), lane 6:C19(EHEC), Lane 7: 46A (EPEC), lane 8: 73A(ETEC), lane 9:C24(EIEC), lane 10:C12 (STEC), lane 11: Non photogenic E. coli, lane 12: negative control (ddwater).

**Figure 9:**
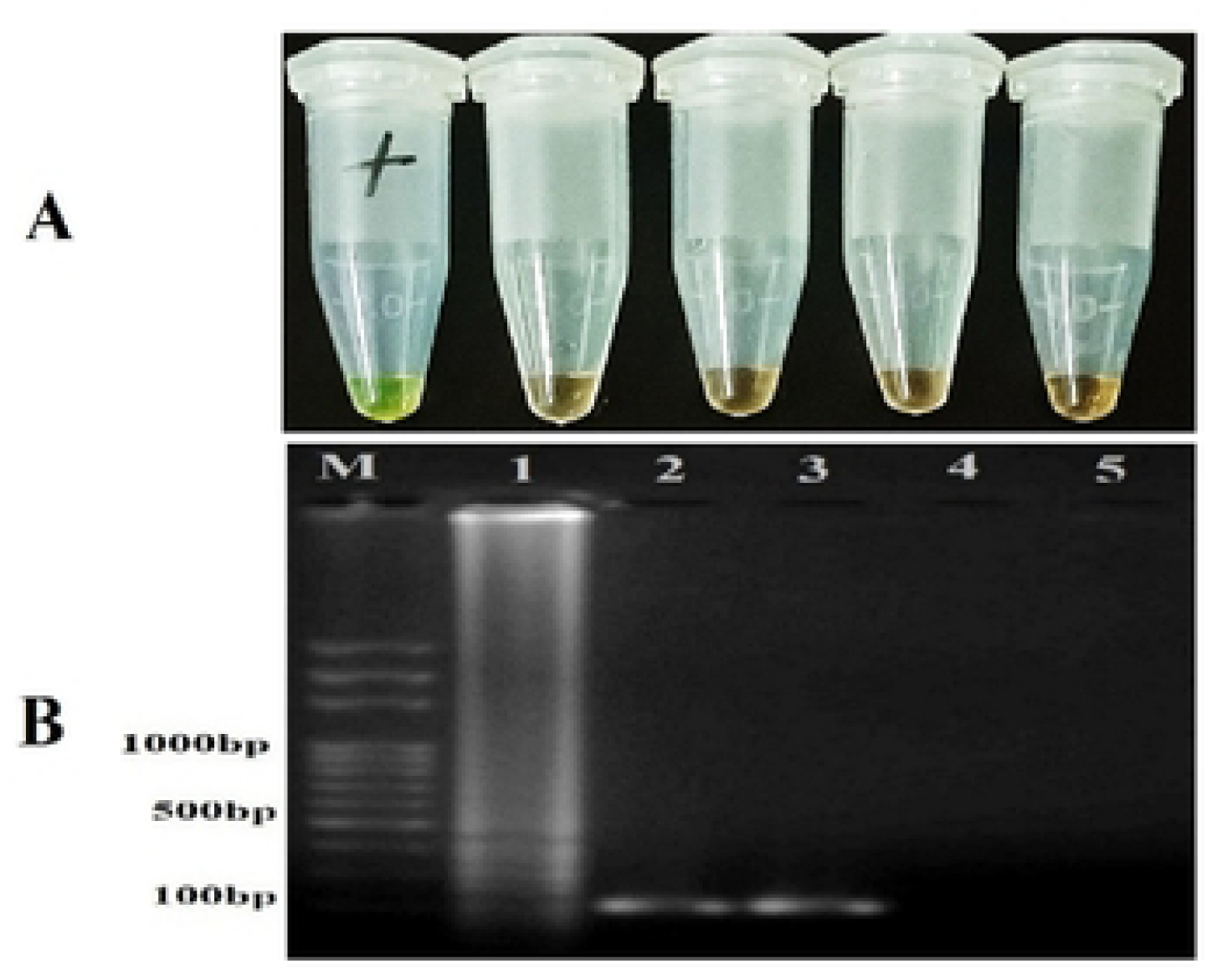
Specificity on non-*E. coli* bacteria: (A) SYBR green I detection; Tube 1:024 (EAEC), Tube 2: klebsiella pneumonia, Tube 3: pseudomonas, Tube 4: Staphylococcus aureus, Tube 5: Salmonella, B) Gel electrophoresis result; lane M: DNA ladder (100plus), lane1:02A, lane2: P1, lane 3: K3, lane 4: S2: lane5: Sal

**Table 5:**
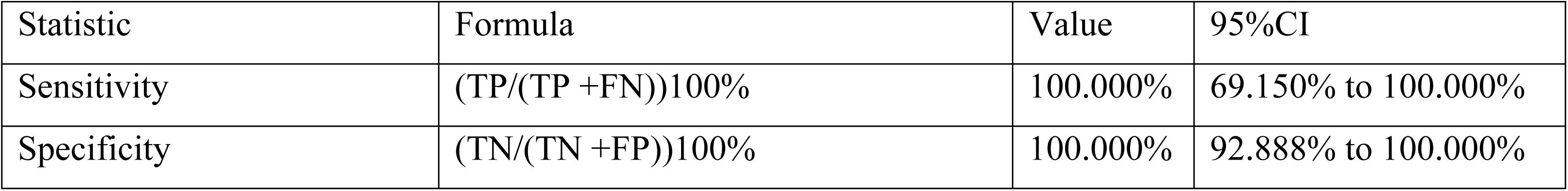
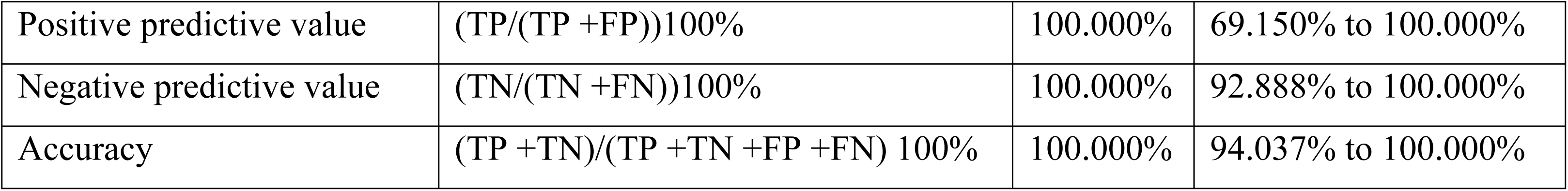
Statistical analysis for evaluating diagnostic performance of the LAMP diagnostic assay using MedCalc Version 23.2.1 Software.

#### 3.3.2. Sensitivity of the LAMP assay

The established LAMP assay achieved 100% sensitivity, confirming the assay’s effectiveness for detecting EAEC without yielding any false-negative results. Furthermore, the study demonstrated a positive predictive value of 100%, indicating that every sample tested positive by the LAMP assay was identified as an EAEC isolate. In parallel, a 100% of negative predictive value was observed, confirming that all negative results accurately indicate the absence of EAEC. These findings show the perfect agreement between the LAMP assay results and the confirmed status of all tested samples, indicating the assay’s high level of diagnostic accuracy and reliability (Table 5).

#### 3.3.3. Cohen’s kappa test statistics

Cohen’s Kappa test statistic(K) was used to evaluate the agreement between the LAMP and PCR assay (Table 6).

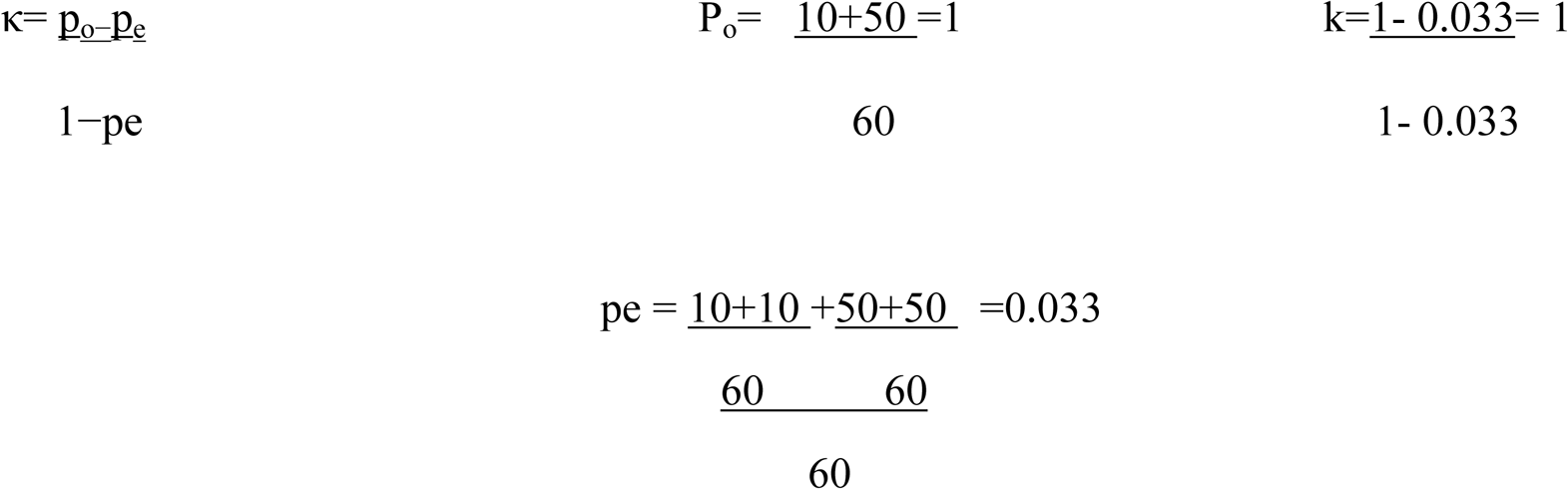

K=1 Shows a perfect agreement between PCR and LAMP assay.

**Table 6:** Kappa test statistics(K).

| LAMP | PCR |  |  |  | Weighted Kappa | 95%CI |
| --- | --- | --- | --- | --- | --- | --- |
|  | Test | Positive | Negative | Total | 1.0 | 1.00 to 1.00 |
|  | Positive | 10 | 0 | 10 |  |  |
|  | Negative | 0 | 50 | 50 |  |  |
|  | Total | 10 | 50 | 60 |  |  |

#### 3.3.4. Lowest detection limit of LAMP assay (LOD)

The developed lamp assay detection limit was determined by performing a series of 10-fold serial dilutions of EAEC template DNA. The LAMP assay showed a positive result at a DNA concentration of as low as 0.098pg of DNA template per reaction, indicating the lowest detection limit of the LAMP assay was 0.098pg /reaction. In contrast, the conventional PCR has a lowest detection limit of 0.98 pg/reaction. The result of both LAMP and PCR across the dilution series are presented in Table 7, and the corresponding amplification results are shown in Figure 10.

**Figure 10:**
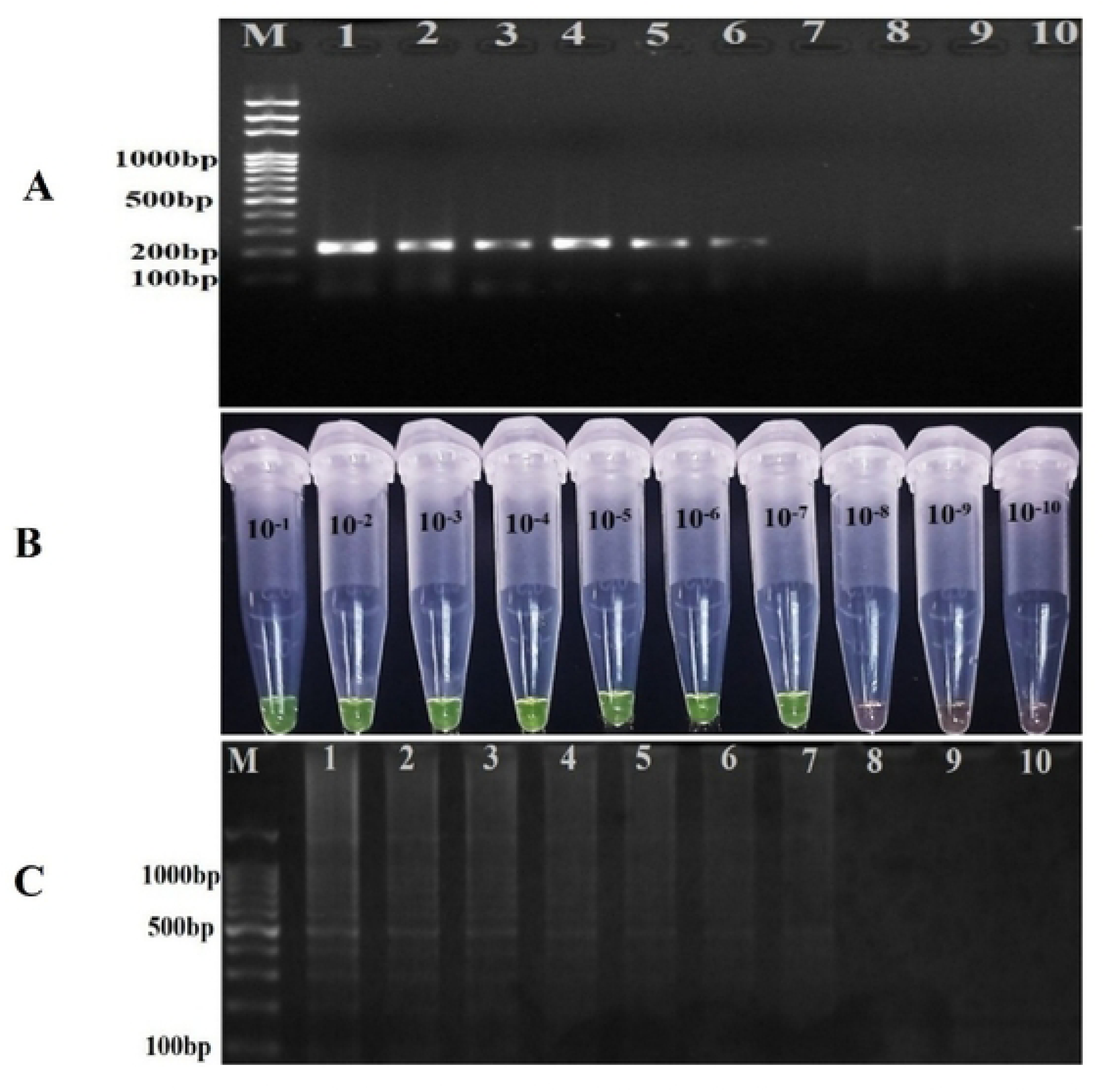
Comparison of sensitivity (serial dilution) of LAMP assay with PCR in detection of EAEC: (A) Electrophoretic analysis of EAEC by polymerase chain reaction (PCR)–amplified products. Lane M: DNA ladder, lane 1: 10^−1^, lane 2: 10 ^−2^, lane 3: 10 ^−3^, lane 4: 10^−4^, lane 5: 10 ^−5^, lane 6:10 ^−6^, lane 7: 10 ^−7^, lane 8: 10 ^−8^, lane 9: 10-^9^lane 10: 10^−10,^ Lane NC: negative control. (B) Sensitivity of LAMP assay detection by SYBR green I: Tube 1: 10^−1^, tube 2: 10 ^−2^, tube 3: 10 ^−3^, tube 4: 10^−4^, tube 5: 10 ^−5^, tube 6:10 ^−6^, tube 7: 10 ^−7^, tube 8: 10 ^−8^, tube 9: 10-^9^, tube10: 10^−10^. (C) Sensitivity of LAMP assay detection analyzed by gel electrophoresis. Lane M: DNA ladder, lane 1: 10^−1^, lane 2: 10 ^−2^, lane 3: 10 ^−3^, lane 4: 10^−4^, lane 5: 10 ^−5^, lane 6:10 ^−6^, lane 7: 10 ^−7^, lane 8: 10 ^−8^, lane 9: 10-9, lane 10: 10^−10^, Lane NC: negative control.

**Table 7:** Summary of serial dilution result, showing a comparative detection limit for both LAMP and PCR.

| Dilution Factor | DNA concentration | DNA concentration per reaction (2 μL template). | LAMP | PCR |
| --- | --- | --- | --- | --- |
| $10^{-1}$ | 49.0ng/ μL | $98 \times 10^1$ ng/reaction | + | + |
| $10^{-2}$ | 4.90ng/ μL | $9.80 \times 10^0$ ng/reaction | + | + |
| $10^{-3}$ | 0.490ng/ μL | $9.80 \times 10^{-1}$ ng/reaction | + | + |
| $10^{-4}$ | 0.0490ng/ μL | $9.80 \times 10^{-2}$ ng/ reaction | + | + |
| $10^{-5}$ | 4.90 pg /μL | 9.8 pg/reaction | + | + |
| $10^{-6}$ | 0. 490 pg /μL | 0.98 pg/reaction | + | + |
| $10^{-7}$ | 0.0490pg/μL | 0.098pg/reaction | + | - |
| $10^{-8}$ | 0.00490pg/μL | 0.0098 pg/reaction | - | - |
| $10^{-9}$ | 0.000490pg/μL | 0.00098 pg/reaction | - | - |
| $10^{-10}$ | 0.0000490pg/μL | 0.000098 pg/reaction | - | - |

Standard plate counting method was used to determine the numbers of bacterial cells detected by the developed LAMP assay. The total viable count of undiluted culture was 8.1×10^9^CFU/mL by using plate-counting method. The results of LAMP and PCR assay showed a notable difference in detection limits. Specifically, the LAMP assay exhibited significantly higher sensitivity, with a detection limit of 81 CFU/mL, compared to 8.1 × 10^2^CFU/mL for conventional PCR. This indicates that the developed LAMP assay is approximately 10 times more sensitive than PCR in identifying the target EAEC in pure culture. Furthermore, in spiked stool sample the lowest detection limit for LAMP was 8.2 × 10^2^ cfu/ g stool. In contrast, 8.2 × 10^4^ cfu/g stool for PCR, indicating that LAMP is 100 times more sensitive than PCR.

## 4. Discussion

Diarrheal disease is a significant cause of death and economic losses in developing countries, it is caused by a diverse group of bacteria, viruses, and parasites. Among bacterial pathogens, diarrheagenic *Escherichia coli (*DEC) are common causes and classified into; Enteroaggregative *Escherichia coli* (EAEC), Enterohemorrhagic *Escherichia coli* (EHEC), enteropathogenic *Escherichia coli* (EPEC), enterotoxigenic *Escherichia coli* (ETEC), enteroinvasive *Escherichia coli* (EIEC), diffusely adherent *Escherichia coli* (DAEC),Shiga toxin producing *Escherichia coli* (STEC),adherent-invasive *Escherichia coli* (AIEC) and cell-detaching *Escherichia coli* (CDEC) strains [1, 26–30]. Enteroaggregative *Escherichia coli* (EAEC) is a significant pathogen responsible for causing acute and persistent diarrhea globally [6, 31].

The widespread prevalence of diarrheal diseases globally emphasizes the need for rapid, affordable, accurate, and easy diagnostic method for the detection of the causative microorganisms. Loop mediated isothermal amplification (LAMP) is an innovative molecular assay emerged as an alternative to PCR for rapid, accurate, and affordable diagnosis. Several studies that have been conducted with LAMP shows its impressive role in addressing the gap in molecular diagnostics[14, 18, 32]. Recently, LAMP assays have been developed for various diarrheal pathogens including pathogenic *Escherichia coli* strains such as Shiga toxin producing *Escherichia coli*, enterohemorrhagic *Escherichia coli*, enterotoxigenic *Escherichia coli[33–36]*. Furthermore, LAMP has been effectively used for the diagnosis of other diarrheal microorganism including, C*lostridioides difficile*, *shigella flexneri, Vibrio cholerae, Entamoeba histolytica*, *rotavirus* and Giardia *lamblia [37–42]*.

In this study we developed LAMP assay for a rapid, sensitive and affordable detection of Enteroaggregative *Escherichia coli* (EAEC). The aaic gene was used as a target gene for LAMP primer designing. The specificity of the developed LAMP assay was evaluated to confirm the assay’s ability for distinguishing EAEC from other closely resemble microorganisms. The present study showed both sensitivity and specificity 100% for EAEC detection, indicating a perfect accuracy for identifying true positive and true negative samples In comparison to previous study where the LAMP assay shows 100% sensitivity and 97.05% specificity for detecting enterohemorrhagic Escherichia coli (EHEC)[36],the specificity of developed lamp assay is higher. The developed LAMP assay and PCR agreement was measured by Cohen’s kappa statistic (k = 1) and the results shows a perfect agreement between PCR and LAMP. This level of agreement is higher than previously reported LAMP assays for EHEC detection [36].

The developed LAMP assay was able to detect DNA at concentrations as low as 0.098 pg /reaction, whereas conventional PCR showed a lower detection limit of 0.98 pg/reaction. This indicates that the developed LAMP assay is approximately 10 times more sensitive than PCR. The results are consistent with previously reported LAMP assay that showed a detection limit of 0.5-0.1pg/reaction for Shiga toxin producing *Escherichia coli* (STEC) strains [43]. In terms of viable bacterial cell, the developed LAMP assay exhibited a 10-fold greater sensitivity than conventional PCR, detecting as low as 81 CFU/mL of EAEC in pure culture, compared to 8.1 × 10^2^ CFU/mL by PCR. Similarly, in spiked stool samples, LAMP detected 8.1 × 10² CFU/g; whereas, PCR required 8.1 × 10⁴ CFU/g. The difference in sensitivity between fecal sample and pure culture maybe related to the presence of inhibitors, such as bile salts and plant-based polysaccharides commonly found in fecal samples[44]. A study on patients with diarrhea reported a bacterial count in diarrheal samples around 2.1 × 10⁹ CFU/g, versus healthy controls averaging ∼5.6 × 10¹⁰ CFU**/**g[45]. Among diarrheal cases where EAEC was the pathogen detected, the bacterial load was around 2.78 × 10⁶ bacteria/g[46]. Therefore, the developed LAMP assay can be used for direct patient diagnosis, as it enables a sensitive detection of EAEC from stool sample, with a lower detection limit of 8× 10² CFU/g. Regarding cost and time, LAMP offer a significant advantage over conventional PCR. LAMP operates under isothermal conditions to produce the results within 60 minutes. This is considerably faster than the 1.5 to 3 hours required for PCR assays. In terms of cost LAMP is more cost effective than PCR. The minimal equipment and simplified protocols make LAMP especially suitable for use in low-resource clinical settings[47].

## 5. Conclusion

The developed LAMP assay has a capability to address the shortage of efficient diagnostic tools in developing countries and significantly reduce both time and cost associated with EAEC diagnosis. The promising performance of the LAMP assay makes it as a robust alternative to PCR for accurate, rapid and affordable detection of EAEC infections, contributing to effective diarrheal disease management and improved public health outcomes.

## Data availability

All data generated and analyzed in this study are available within the manuscript.

## Acknowledgments

The authors would like to thank institute of biotechnology, Addis Ababa university for their kind cooperation in facilitating this study.

## Abbreviation

EAEC: Enteroaggregative *Escherichia coli*
HEp-2: human epithelial cell
TP: True positive
TN: True negative
NPV: Negative predictive value
PPV: positive predictive value

